# Isolation and Identification of taxonomically diverse bacterial endophytes from citrus in Punjab Pakistan

**DOI:** 10.1101/2022.01.04.474905

**Authors:** Sehrish Mushtaq, Muhammad Shafiq, Tehseen Ashraf, Fahim Qureshi, Muhammad Saleem Haider, Sagheer Atta

**Author notes:** These authors contributed equally to this work (Sehrish Mushtaq, Sagheer Atta). These authors also contributed equally to this work.

## Abstract

Citrus is an economically important fruit crop grown in all provinces of Pakistan, while Punjab accounting for 95 percent of total production due to its favorable climate for citrus production. Commercially grown varieties in Pakistan include sweet oranges, grapefruits, Mandarine, Lime, and lemon. The goal of this research was to see how diverse the cultivable bacterial populations are found in citrus cultivars. Out of 90 isolated cultures, 37 endophytic bacterial species and 15 different genera of bacteria were characterized based on morphological, biochemical, and molecular methods from citrus leaves. All the isolated bacteria were subjected to PCR amplification through 16S rDNA followed by sequencing. RDP base classification revealed that class Bacilli has the largest percentage of isolates, whereas class Alpha, Beta, and Gamma Proteobacteria have the lowest percentage among all genotypes used. According to the findings, the phylum Firmicutes contains a common genus (Brevibacterium, 1%; Enterococcus, 6%; Staphylococcus, 7% and Bacillus, 60%). Alpha (Rhizobium) beta (*Burkholderia cepacia*; *Comamonas terigena*) gamma Proteobacteria (*Enterobacter hermachei* (1%), *Klebsiella pneumoniae* (1%), *Proteus mirabilis* (8%), Pseudomonas *aeruginosa* (5%), *Psychrobacter pulmonis* and *Yersinia molalretti* (1%) respectively. These results revealed that cultivars of the plants might contribute to the structure and endophytic bacterial communities associated with citrus. Endophytes extracted from leaf samples of different citrus cultivars in Pakistan are reported for the first time. The idea of employing endophytes bacteria to produce enzymes stimulate plant growth, and its purpose as a biological control agent will be investigated in the future.

## Introduction

Citrus has traditionally been recognized as one of the most popular fruit, as it is enriched with minerals, vitamins A, B, and C, ascorbic acid, and also possesses strong antioxidant potential [1, 2]. Pakistan is the world’s number one producer of Kinnow mandarin [3]. A widespread kind of microorganisms exists in the phyllosphere (Surface of the plant). Bacteria are the most common microbes found in the phyllosphere. Endophytes (bacteria that live inside plant tissues) and epiphytes (bacteria that live on the surface of plants) are both examples of phyllosphere bacteria [4, 5]. The research on plant-based endophytes is critical for understanding the diverse bacterial interactions that occur in certain environments, which help to improve their biotechnological applications [6, 7]. The endophytes that reside within the surface of the leaf form complex population dynamics that are important for agriculture and the environment. Plant health can be improved or arrested by the bacterial diversity associated with the leaf, which can also drive the colonization and infection of tissues by phytopathogens [8, 9].

Microorganisms that live in a mutualistic connection with a plant are known as endophytes [10, 11]. Plant processes such as growth promotion, nutrient uptake, abiotic stress tolerance, and pathogen infection inhibition are thought to be supported by them [6]. Bacterial endophytes (BE) are found in different parts of plants such as roots, stems, and leaves [12, 13]. The population density of BE is affected by multiple factors, including the plant’s developmental phase [7, 14], cultivar (genotype) [14, 15], the portion under examination [16], comprising the type of plant being studied, and the interaction between microbes and ecological factors [17]. The population density of BE could be 10^2^ to 10^9^ people per square kilometer [18]. Commonly, BE present in lower numbers as compared to the rhizospheric bacteria [19]. They are not limited to a single species in a single plant host, but they could be observed in multiple genera and species. Endophytic bacteria have been extracted from different segments of *M. micrantha*, including the roots, lamina, and petiole [20]. In previous studies, endophytes have been isolated from cottonwood (*Populus deltoids*) [21], grapevine (*Viti vinifera*) [22], poplar (*Populus alba*) [23], sweet potato (*Ipomoea batatas*) [24], potato [25, 26], soybean (*Glycine max*) [27] and tomato (*Solanum lycopersicum*) [28].

Pakistan has an agriculturally dominated economy with large areas of fertile land and a diversified geographical region and climate. There has been limited research on microbial biodiversity, and no such type of research on the diversity of BE from Citrus in Pakistan has ever been reported. It is so believed that bacterial species found in Pakistan carrying a wide variety of endophytic bacteria will be investigated to find the real picture. This study will provide information regarding the different strains of bacteria that are residing in citrus leaves and still need to be identified from different geographical locations of the country.

## Methods

### Sample collection and isolation of bacteria

Citrus leaf samples were collected from orchards in the Punjab districts of Lahore, Multan, Mian Chanu, Sahiwal, Faisalabad, and Sargodha. For subsequent processing, leaves from thirty-two different citrus varieties were collected and stored at −80°C. To isolate endophytes, 3-4 cm of citrus leaf midrib part were sanitized with NaOCl solution (0.6%) for three min and then rinsed thrice with sterilized deionized autoclaved water. A homogeneous solution of crushed mid rib portion and deionized water was prepared and streaked on Nutrient agar (NA) medium plates, which were incubated at 28°C for 1-2 days. Purified colonies were culture on NA plates and incubated at 28°C for 1 day until growth appears. Following Bergey’s manual of systemic bacteriology, pure cultures of bacterial isolates were identified using morphological and biochemical methods [29].

### Identification of bacterial isolates by 16S rDNA

Genomic DNA of Bacterial endophytes was extracted by the Cetyl trimethyl ammonium bromide (CTAB) method followed by [30]. Single-cell colonies of bacteria were culture in 5mL of NA for 24 hours and centrifuged at 13000 rpm for 2 min. Then the pallet was dissolved in mixture of TAE buffer, 10% SDS, proteinase k (20 mg/ml) and incubated at 37°C for 60 min. 5M NaCl (100μL) and CTAB (80μL) were added and incubated for 10 minutes at 65°C. After that 750μL of Chloroform Isoamyl Alcohol (24:1) were added and centrifuged for 10 min. About 400μL of the upper phase was removed and shifted to another tube. Then (700μl) of Phenol Chloroform was added and centrifuged for 10 min and repeat this step. After that (20μL) of 3M Sodium Acetate and (500μL) of Absolute Ethanol were gently mixed and kept at −20°C for 12 h. After that tubes were spin (centrifuged) again at 13000 rpm for 10 min and the supernatant was wasted. The DNA pellet was washed with 70% of ethanol and dissolved in distilled autoclave water. Extracted DNA was confirmed on 1% agarose gels containing EtBr (ethidium bromide) (0.5 μg/ mL) and visualize under UV light.

Genomic DNA of 90 bacterial isolates was subjected to PCR by 16SrRNA primers 27-F (5’ AGAGTTTGATCMTGGCTCAG 3’), 1492-R (5’ ACCTTGTTACGACTT 3’), and following PCR conditions reported by (Trivedi et al. 2011). PCR products were Gel purified and sent for Sequencing to Macrogen South Korea. The obtained sequences were aligned with the reported ones in the Gene Bank using BLAST (Basic Local Alignment Search tool) at the NCBI (National Centre for Biotech Information). Apart from NCBI Genbank, the isolated bacterial sequences were classified using the Ribosomal Database Project (RDP Hierarchy Browser). As previously stated, all the obtained sequences of bacterial endophytes after proper identification were submitted to NCBI Gene bank Database and attained their Accession numbers (Table 3).

### Evolutionary Study of Bacterial strains

After sequence analysis, MEGA 7.0 software was used to construct a phylogenetic evolutionary tree through multiple sequence alignment of 16S rDNA gene sequences with other reported ones [31]. A bootstrap analysis of 1000 replications was used to calculate a confidence value for the aligned sequence data set. To investigate the evolutionary relationship between bacterial endophytes, a phylogenetic tree was created using the neighbor-joining algorithm. The signature sequence of the endophytes was also identified through MEGA 6.0 (Muscle) software following the alignment of isolated strains with reported strains (NCBI Data Base). These signatures help in determining the sequence that is unique to a given genus or species when compared to other reported genera or species.

## RESULTS

This study provided a brief overview of the distribution of leaf endophytic bacteria found in citrus varieties, contributing significantly to the study of microbial populations in Pakistan’s environment.

### Morphological and Biochemical Identification of extracted endophytic bacterial strains

Extraction of bacteria from leaf midribs was accomplished using mince soaked techniques on NA medium, and identification based on colony morphology (shape, texture, color, elevations, margins and, transparency of colony) was accomplished using Bergey’s Manual of Systematic Bacteriology Smibert and Krieg (1981), and findings have been summarized in (Table 1 and 2).

**Table 1:**
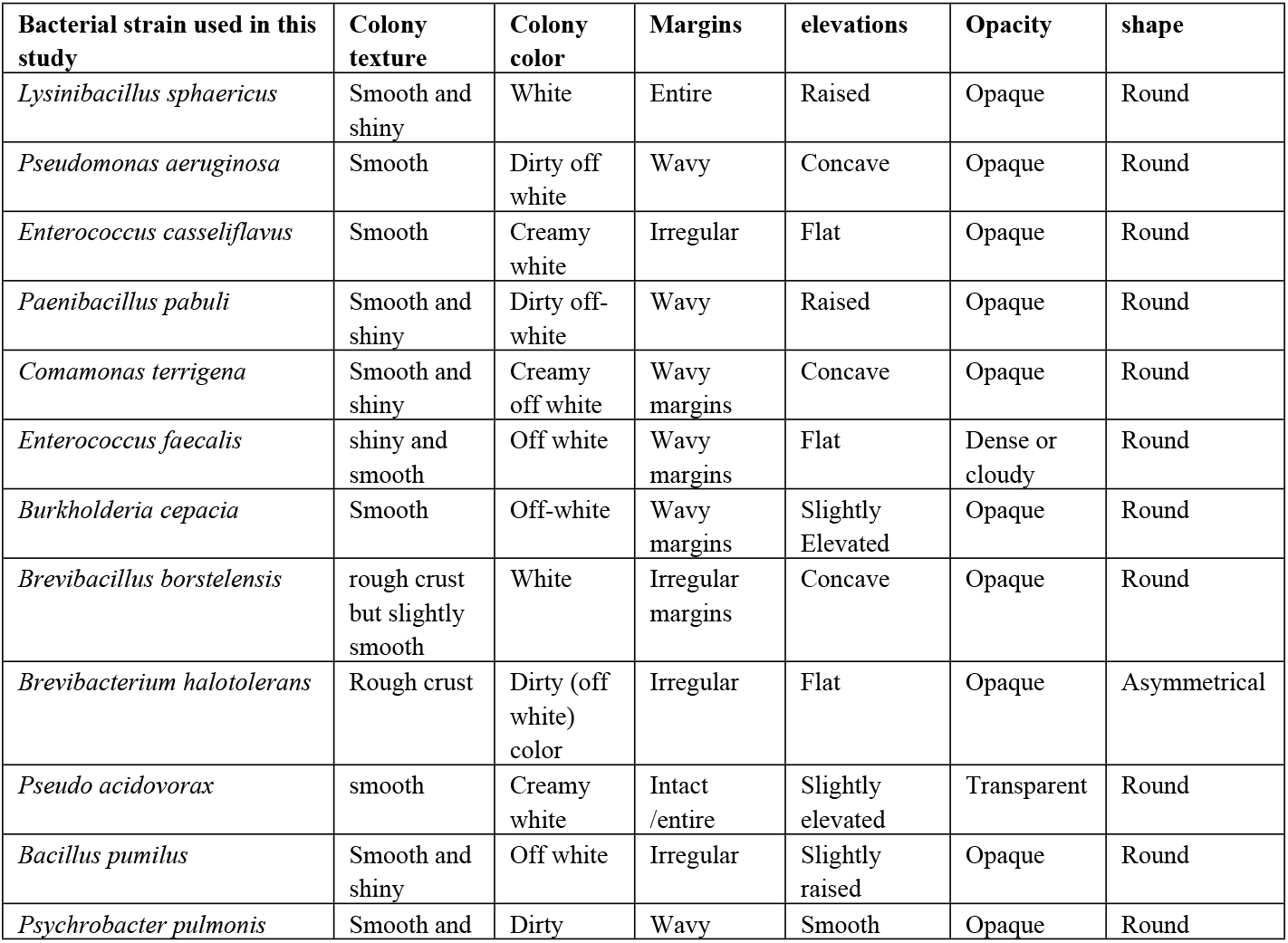
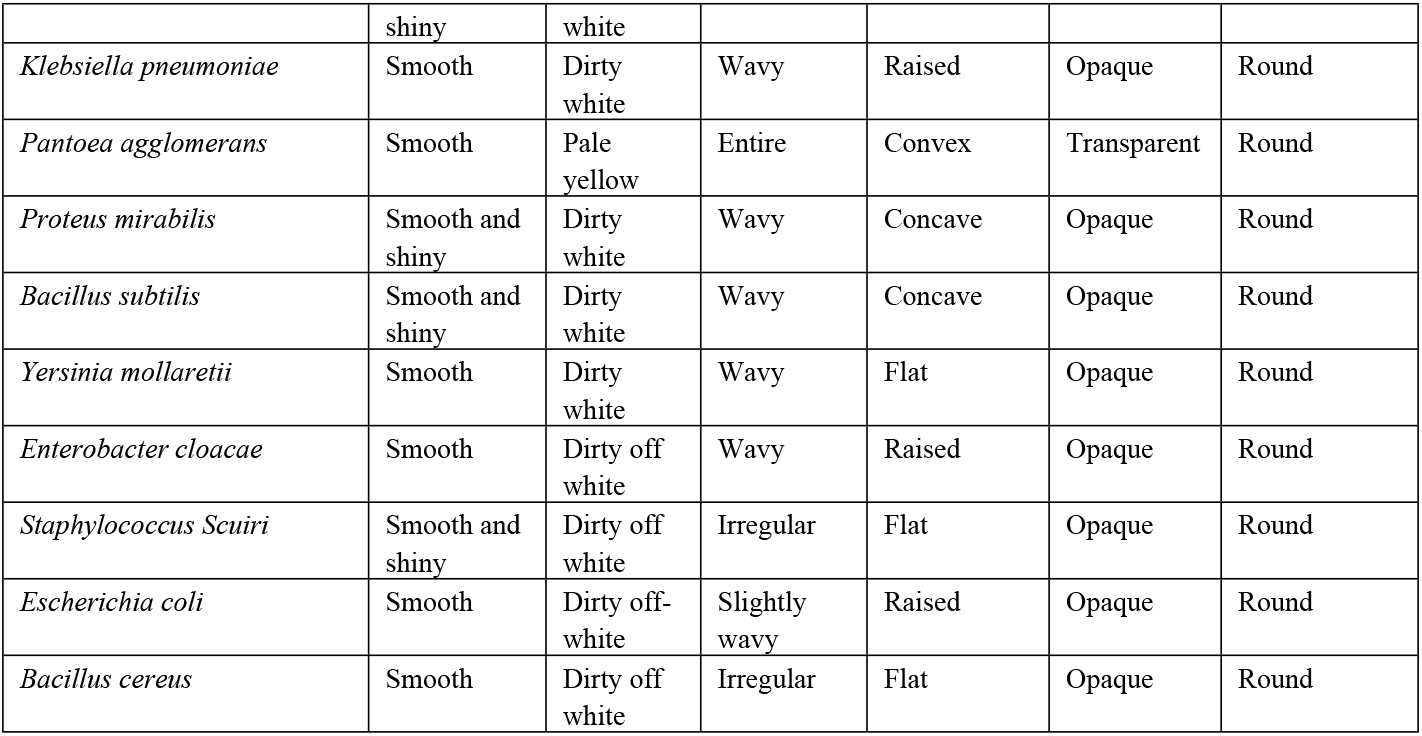
Morphology based determination of bacterial endophytes from Leaf tissue of various citrus genotypes.

**Table 2:** Biochemical assessment of endophytes extracted from leaf mid rib part of different citrus cultivars.

| Identified Bacterial Strains | Gram type | Cell shape | Staining of Capsule | Motility test | Spore type | Methyl red test | Citrate utilization test | Indole test | Hydrogen sulphide test | Oxidase test | Catalase test | Nitrate reduction test | Growth at 40 °C | Growth at 25°C | Growth at 2% NaCl |
| --- | --- | --- | --- | --- | --- | --- | --- | --- | --- | --- | --- | --- | --- | --- | --- |
| *Brevibacillus borstelensis* | Pos. | Rod | Neg. | Pos. | Pos. | Pos. | Neg. | Neg. | Neg. | Neg. | Pos. | Neg. | Neg. | Pos. | Neg. |
| *Comamonas terrigena* | Neg. | Rod | Neg. | Pos. | Neg. | Neg. | Pos. | Neg. | Neg. | Pos. | Pos. | Pos. | Neg. | Neg. | Neg. |
| *Burkholderia cepacia* | Neg. | Cocci | Neg. | Neg. | Neg. | Neg. | Neg. | Neg. | Neg. | Neg. | Neg. | Neg. | Neg. | Neg. | Neg. |
| *Enterococcus casseliflavus* | Neg. | Rods | Neg. | Neg. | Neg. | Neg. | Neg. | Neg. | Neg. | Neg. | Pos. | Neg. | Neg. | Neg. | Neg. |
| *Paenibacillus pabuli* | Pos. | Rod | Neg. | Pos. | Pos. | Pos. | Neg. | Neg. | Neg. | Neg. | Pos. | Neg. | Neg. | Neg. | Neg. |
| *Enterococcus faecalis* | Pos. | Cocci | Neg. | Neg. | Neg. | Neg. | Neg. | Neg. | Neg. | Neg. | Neg. | Pos. | Neg. | Neg. | Neg. |
| *Lysinibacillus sp.* | Pos. | Rod | Neg. | Pos. | Pos. | Neg. | Pos. | Neg. | Neg. | Pos. | Pos. | Neg. | Neg. | Pos. | Neg. |
| *Enterococcus faecalis* | Neg. | Rods | Neg. | Pos. | Neg. | Neg. | Pos. | Neg. | Neg. | Neg. | Neg. | Pos. | Neg. | Neg. | Neg. |
| *Brevibacterium halotolerans* | Pos. | Rod | Neg. | Neg. | Neg. | Pos. | Neg. | Neg. | Neg. | Neg. | Pos. | Neg. | Neg. | Neg. | Neg. |
| *Enterobacter cloacae* | Neg. | Rod | Neg. | Pos. | Neg. | Neg. | Pos. | Neg. | Pos. | Pos. | Pos. | Neg. | Neg. | Neg. | Neg. |
| *Escherichia coli* | Neg. | Rod | Neg. | Pos. | Neg. | Pos. | Neg. | Pos. | Neg. | Neg. | Pos. | Pos. | Neg. | Neg. | Neg. |
| *Staphylococcus haemolyticus* | Pos. | Cocci | Neg. | Neg. | Neg. | Pos. | Pos. | Neg. | Neg. | Neg. | Pos. | Pos. | Neg. | Pos. | Neg. |
| *Yersinia mollaretii* | Neg. | Rods | Neg. | Pos. | Neg. | Pos. | Neg. | Neg. | Neg. | Neg. | Pos. | Neg. | Neg. | Pos. | Neg. |
| *Bacillus pumilus* | Neg. | Rods | Neg. | Pos. | Neg. | Neg. | Pos. | Neg. | Neg. | Neg. | Pos. | Pos. | Neg. | Neg. | Neg. |
| *Pseudomonas aeruginosa* | Neg. | Rod | Neg. | Pos. | Neg. | Neg. | Pos. | Neg. | Neg. | Pos. | Pos. | Pos. | Neg. | Neg. | Pos. |
| *Klebsiella sp.* | Neg. | Rod | Pos. | Neg. | Neg. | Neg. | Pos. | Neg. | Neg. | Neg. | Pos. | Pos. | Neg. | Pos. | Neg. |
| *Proteus Sp.* | Neg. | Rod | Neg. | Pos. | Neg. | Pos. | Pos. | Neg. | Pos. | Neg. | Pos. | Pos. | Neg. | Neg. | Neg. |
| *Pseudoacidovorax* | Neg. | Cocci | Pos. | Pos. | Neg. | Neg. | Neg. | Neg. | Neg. | Neg. | Neg. | Neg. | Neg. | Pos. | Neg. |
| *Psychrobacter pulmonis* | Neg. | Rods | Neg. | Pos. | Neg. | Pos. | Neg. | Neg. | Neg. | Neg. | Pos. | Neg. | Neg. | Pos. | Neg. |
| *Bacillus cereus* | Neg. | Rods | Neg. | Pos. | Neg. | Neg. | Neg. | Neg. | Neg. | Pos. | Neg. | Neg. | Neg. | Neg. | Pos. |
| *Pantoea Sp.* | Neg. | Rod | Neg. | Pos. | Neg. | Neg. | Pos. | Neg. | Neg. | Neg. | Pos. | Pos. | Neg. | Neg. | Neg. |
| *Bacillus subtilis* | Pos. | Rods | Neg. | Pos. | Pos. | Neg. | Pos. | Neg. | Neg. | Pos. | Pos. | Pos. | Neg. | Pos. | Neg. |

**Note:** Pos. and Neg. Abbreviations represent the positive and Negative respectively.

### Diversity of Bacterial Endophytes

To get a representative sample of the cultivable endophytes living inside leaf tissues (Citrus), Single-cell and morphologically different colonies of bacteria were picked for 16S rDNA sequencing. A BLAST homology search resulted in the identification of 37 bacterial endophytic species. The overall diversity of the citrus endophytes is represented in the form of Pie graphs (Figure 1 & 2) which showed the distribution of bacterial groups and Genus based on their prevalence percentage in samples.

**Fig.1:**
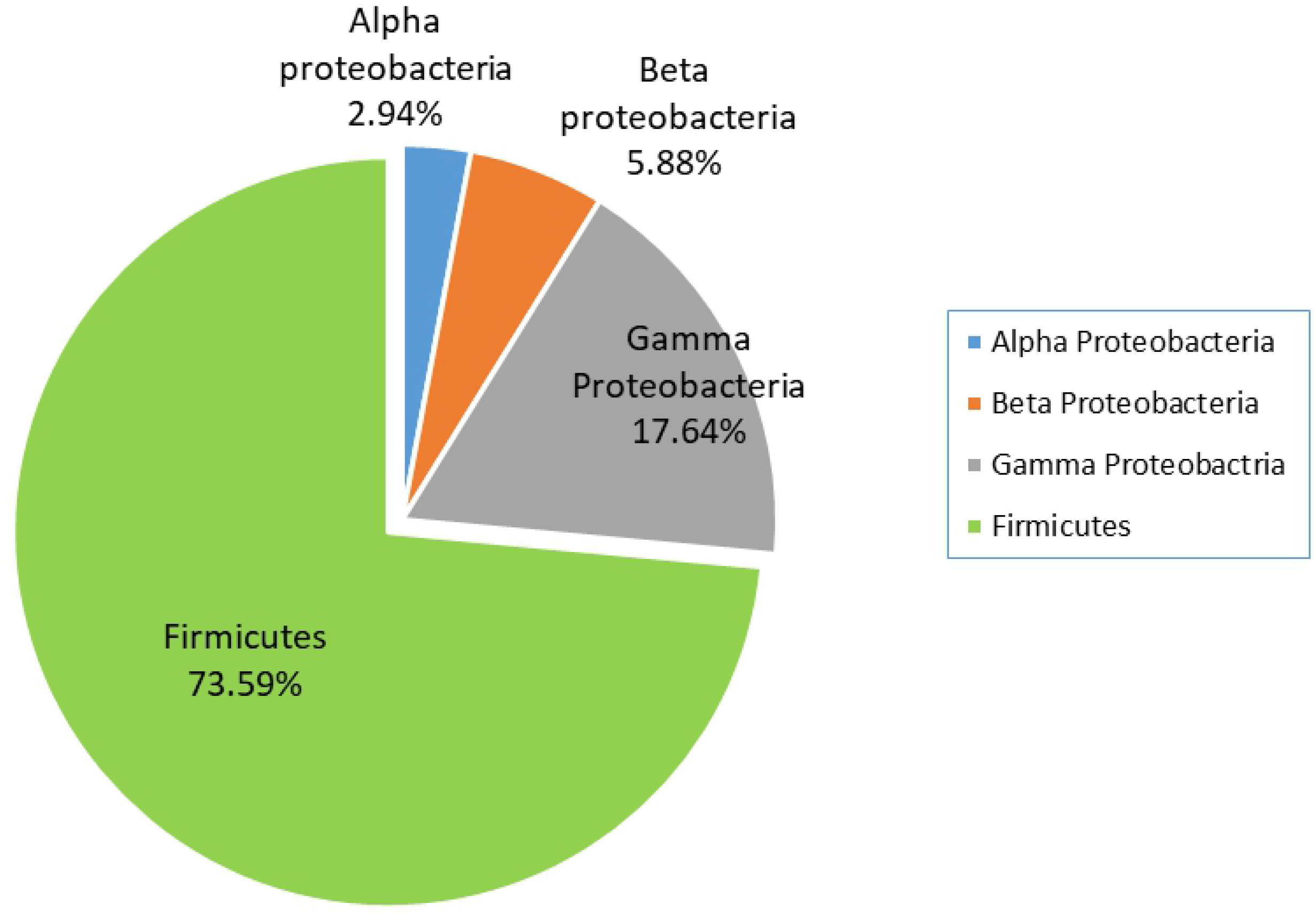
Abundance of endophytes extracted from different citrus genotypes based on Phylum

**Fig.2:**
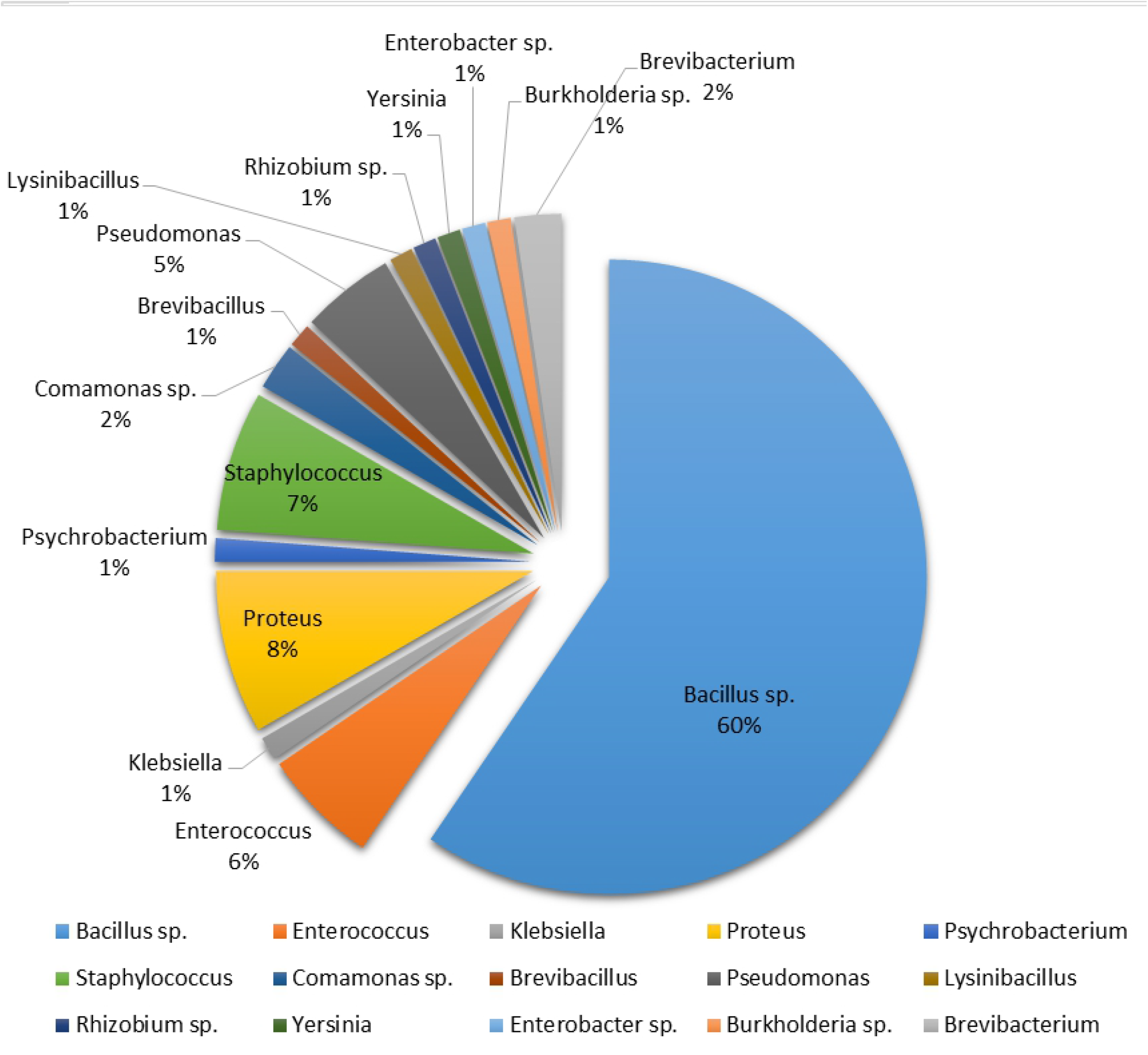
The abundance of endophytic bacteria extracted from various citrus genotypes based on Genus

These bacterial endophytes were classified into two main groups: Firmicutes and Proteobacteria. About 73.59% of Firmicutes were found in citrus samples assessed. Proteobacteria was further comprised of three groups including (2.94%) of Alpha proteobacteria, (5.88%) of beta proteobacteria, and (17.64%) of gamma proteobacteria. The results showed that the majority of genera belonged to the phylum Firmicutes made up of genus (Bacillus accounts for 60%, Enterococcus for 6%, Staphylococcus for 7%, and Brevibacterium for 1%). Additionally, during this study, the phylum Proteobacteria (Alpha (Rhizobium) beta (*Burkholderia cepacia; Comamonas terigena*), gamma Proteobacteria including *Enterobacter hermachei* (1%), *P. aeruginosa* (5%), *Psychrobacter pulmonis* 1%), *P. mirabilis* (8%), *Yersinia molaretti* (1%), and *K. pneumoniae* (1%) were detected. These isolates had a high level of 16S rDNA gene similarity with the genera Bacillus (60%), Staphylococcus (7%), and Proteus (8%), all of which belong to the division Bacilli, while Proteus mirabilis (8%), a member of the Gamma Proteobacteria, displayed the highest level of diversity among all isolates. *Bacillus cereus* remained the frequently discovered and abundant endophyte in citrus leaves extract, according to a research of endophytes prevalence both species and genus levels as illustrated in (Figure 2).

The findings of detection of bacterial species from the leaf of various citrus cultivars and subsequent characterization of partial 16S rRNA gene sequences, such as host, location, and Accession no., are presented in the table below (Table 3).

**Table 3.**
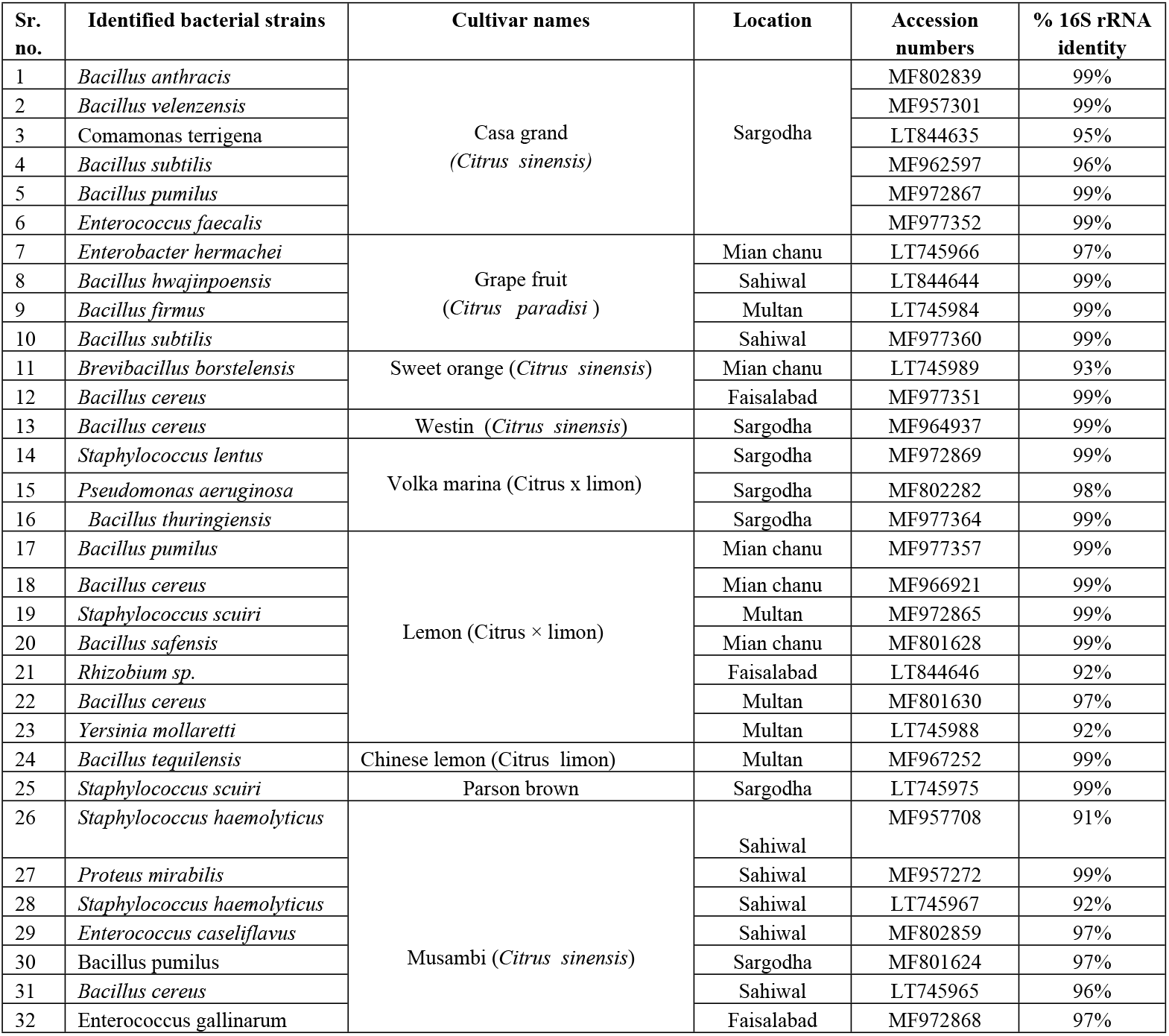
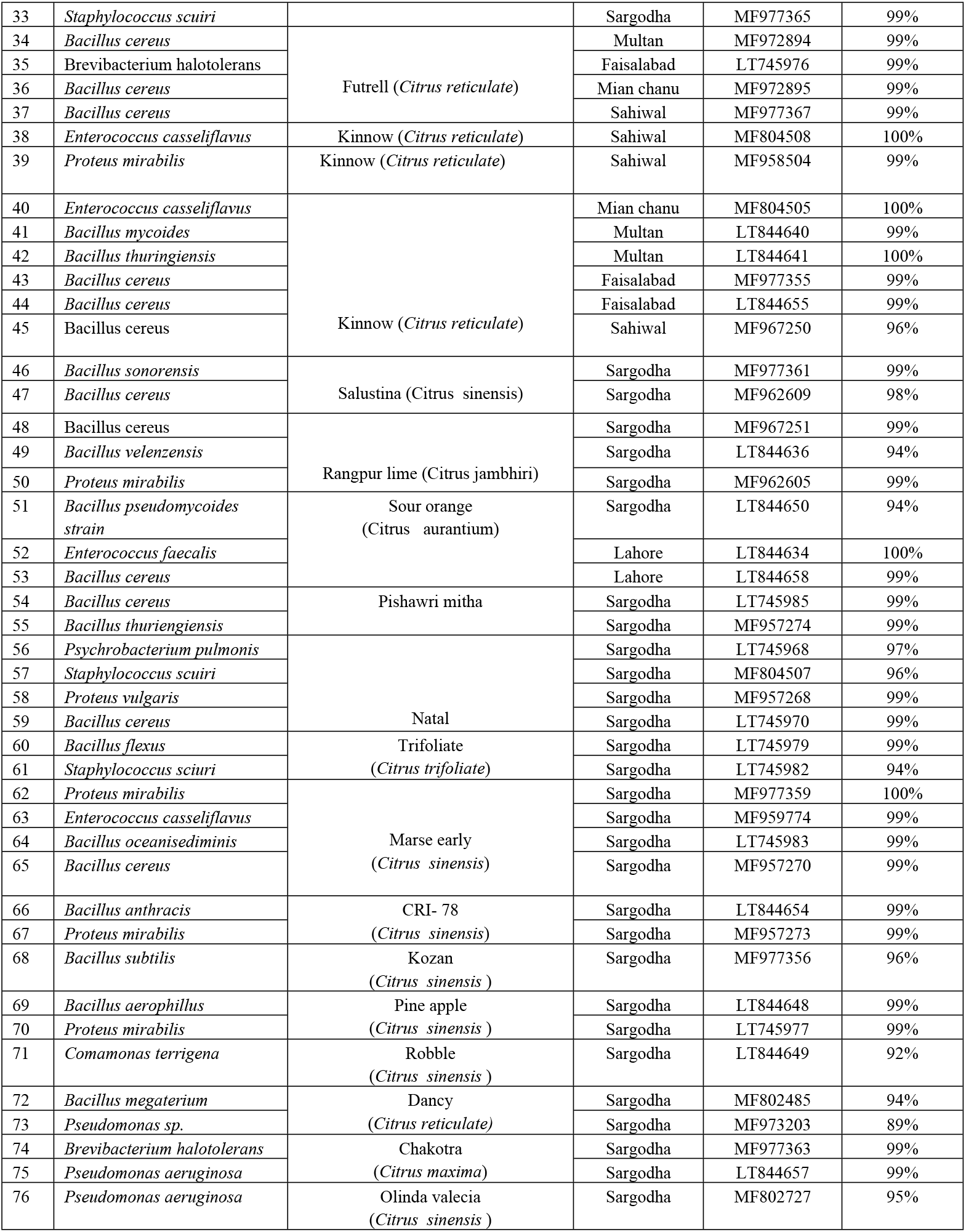
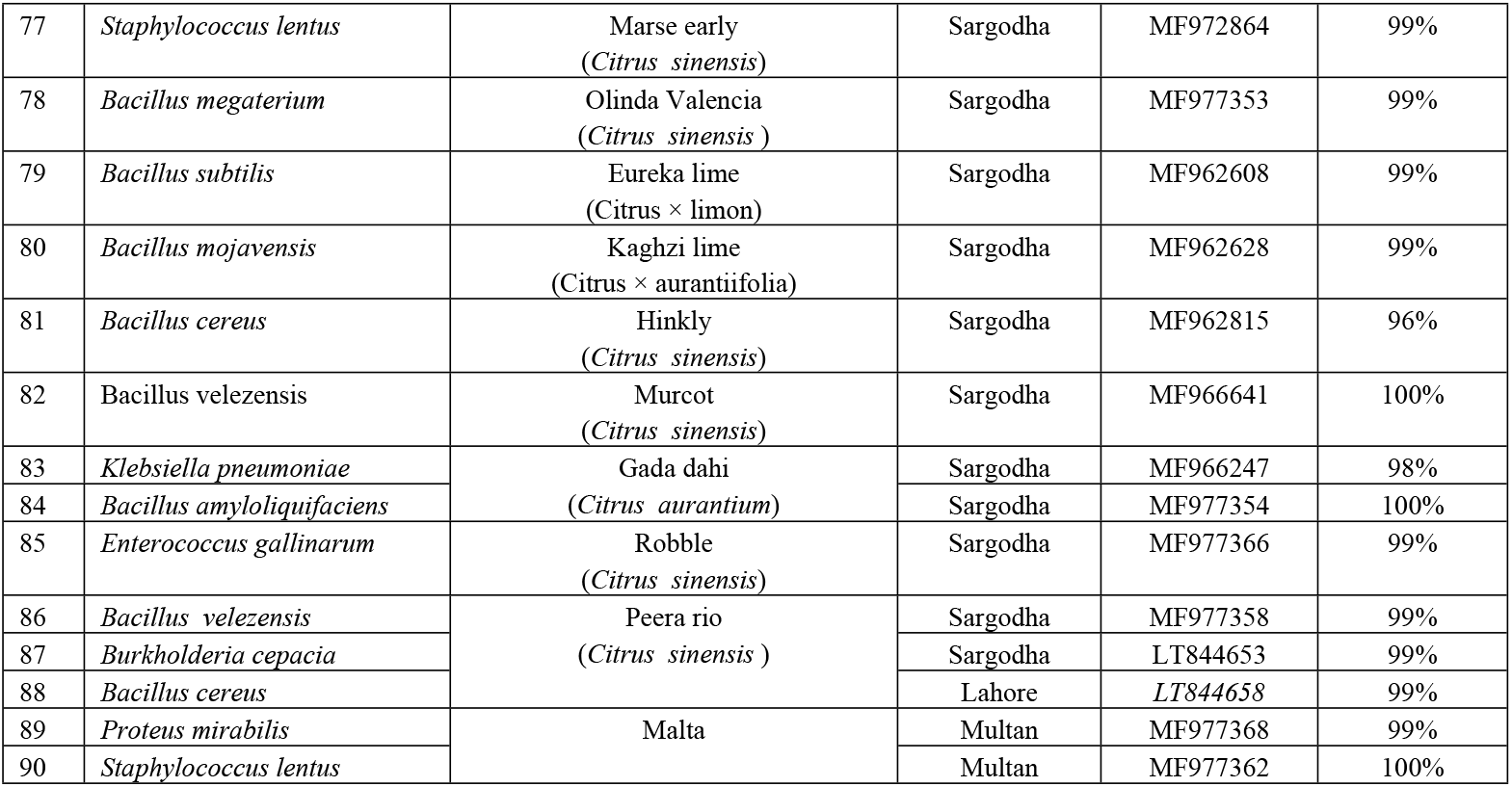
Molecular Characterization based on Partial 16S rRNA sequencing of endophytes extracted from leaf mid rib part of different citrus cultivars.

For phylogenetic investigations, all isolates with a nucleotide sequence identity of 91-100 percent were selected. Using MEGA 6 software and a 1000 bootstrap value, a neighbor-joining dendrogram was contracted. During sequence alignment analysis, 16S rDNA bacterial sequences isolated from different citrus samples were used along with reported species with identities greater than 97%. This phylogenetic analysis contains 90 bacterial strains from the four classes described as (Bacilli, Alpha proteobacteria, Beta proteobacteria, and Gamma proteobacteria) as shown in (Figure 3).

**Fig.3:**
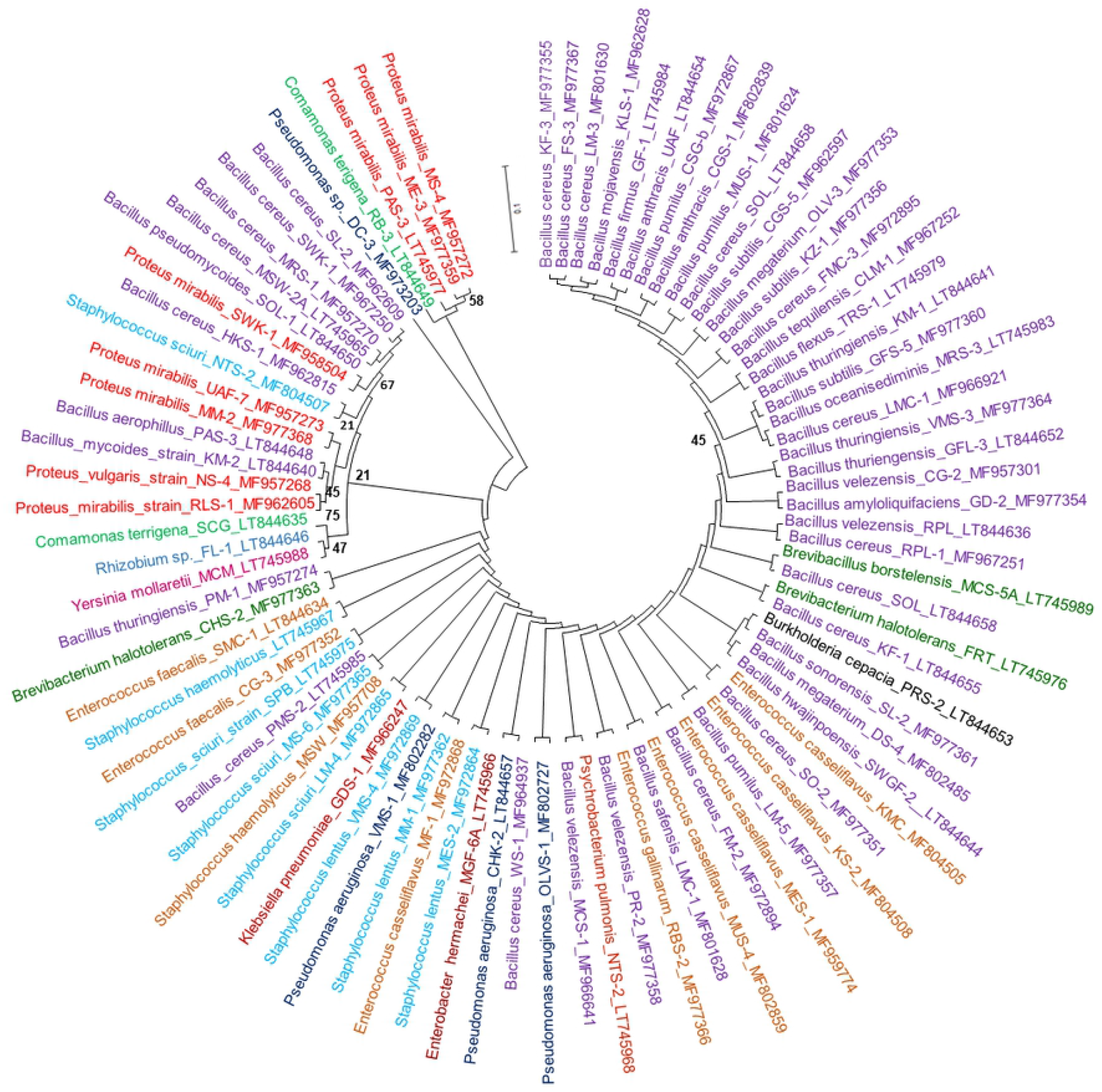
Illustrate the evolutionary relationships of bacterial endophytes obtained from different citrus genotypes. This evolutionary tree contains 22 bacterial endophytes constitute three classes (Bacilli, Beta Proteobacteria, and Gamma Proteobacteria). All of the analysed samples by 16S rDNA were 97 percent identical to known bacterial strains.

### Identification Patterns/Signature Sequences of bacteria

16S ribosomal RNA gene (rRNA) sequencing is considered the gold standard in bacterial identification and classification according to modern taxonomists, with more than 100,000 sequences available in public databases (NCBI). The 16S rRNA gene has conserved regions that can be used to construct PCR primers that can amplify diverse portions of the 16S rRNA gene from bacteria. Hypervariable sections with species-specific signature sequences are included among the fragment, which can be used to identify bacteria up to the species level. All identified bacterial endophytes’ signatures patterns are shown in (Table 4).

**Table 4:**
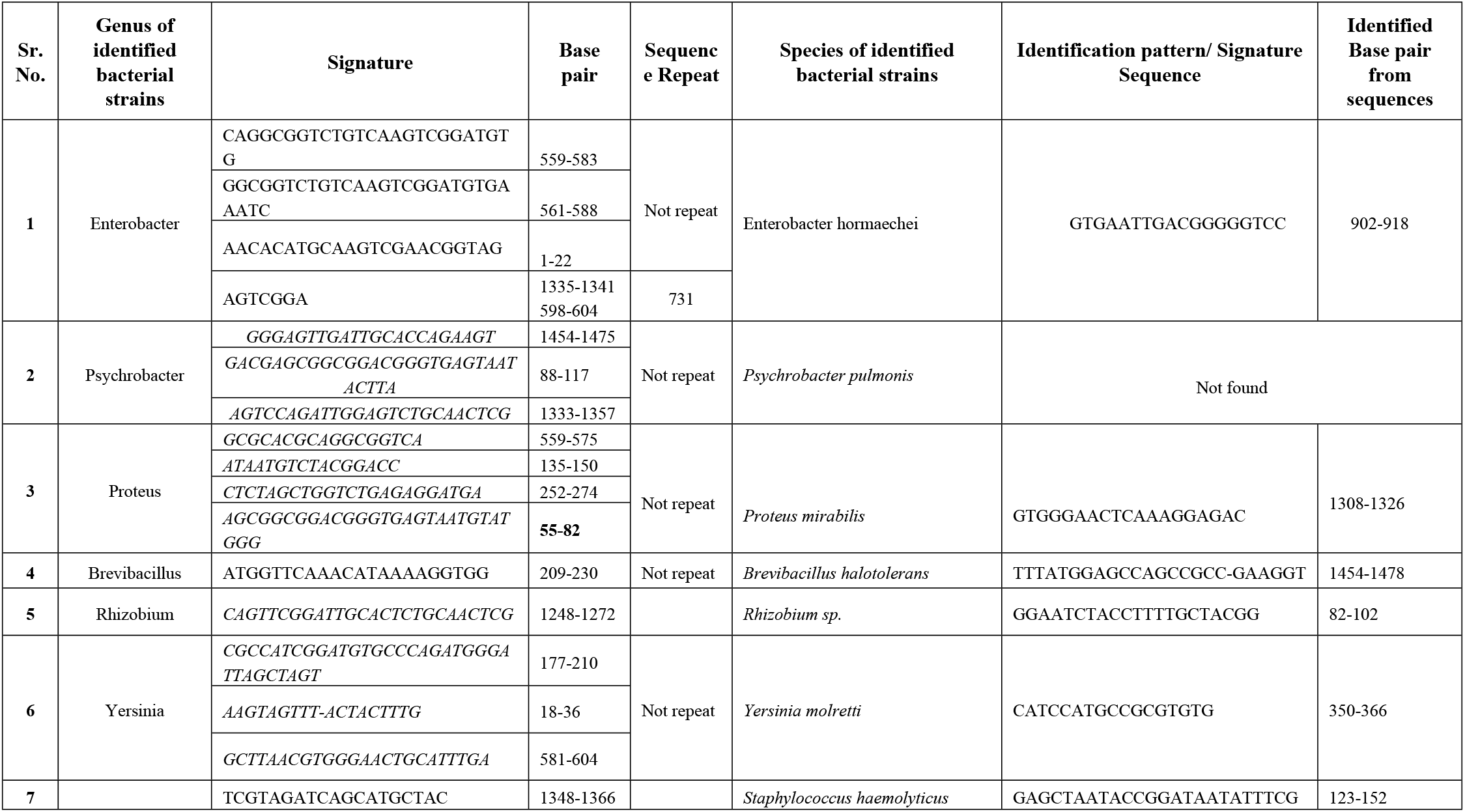
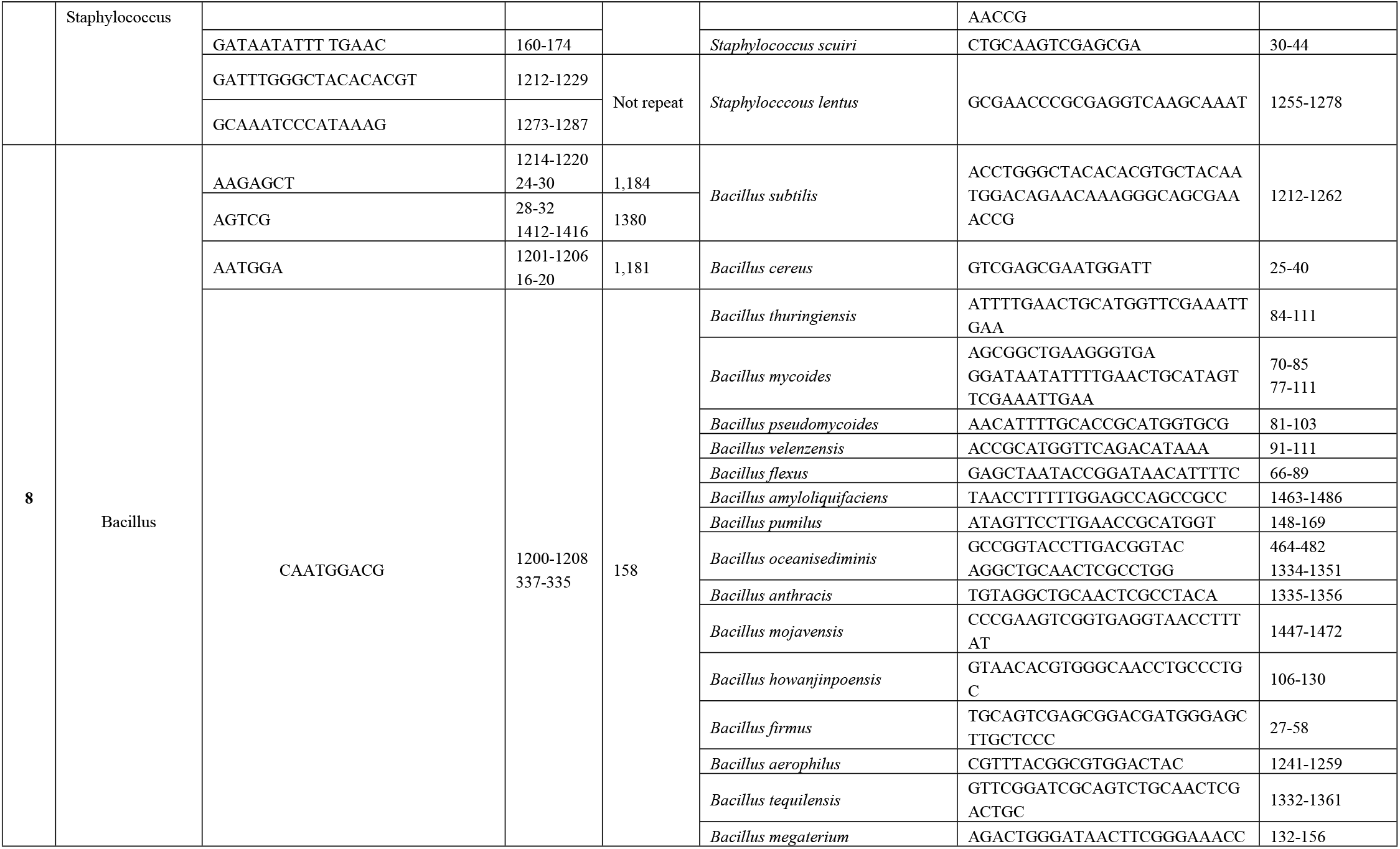
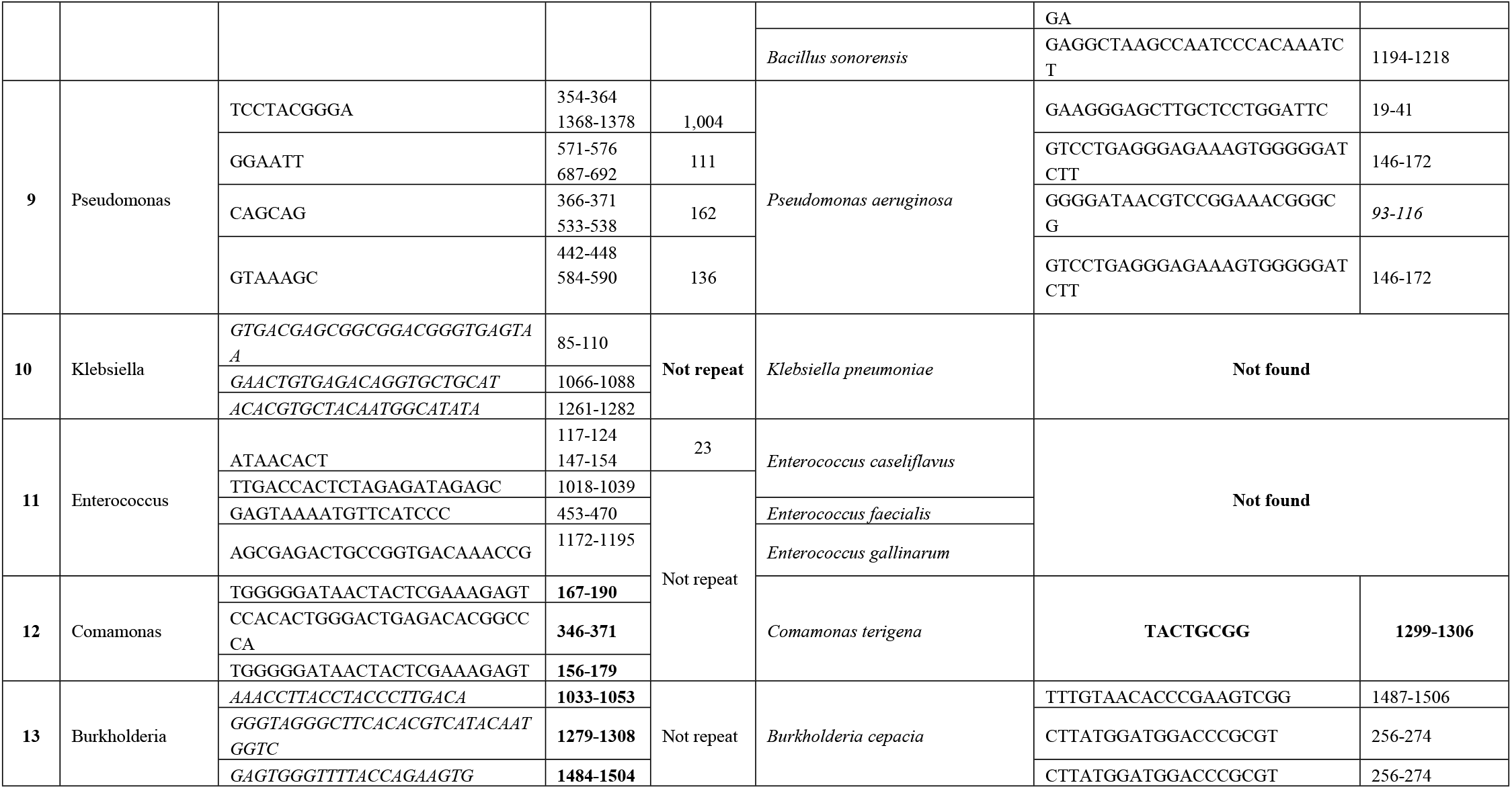
Descriptive analysis of isolated Bacteria at the genus or species level, demonstrating possible identification patterns/signature sequences.

## Discussion

### Diversity of bacteria from Citrus

Microbial community research can help towards a more comprehensive understanding of the complex structure and function of microbial flora, and it may lead to rapid identification of novel bacterial strains [33, 34]. This study provides a thorough examination of the diversity and makeup of microbial communities within the leaf tissue of citrus. Endophytes are important in food safety, agricultural production, and phytoremediation research. The complexity of endophytic microbial flora and the processes that affect their formations in non-cultivated plants, on the other hand, is indeed unexplored. One of the really interesting and relatively unknown fields of agriculture in Pakistan is the exploration of the epiphytic microbial biodiversity of citrus. It has been found that the population of endophytic bacteria within leaf tissue is approximately proportional to the biological influence imparted on the hosts. This is a fascinating theory that needs to be tested on endophytes in different plants. Furthermore, the number of bacterial endophytes inhabiting plant inner parts may be dynamic and depends on a variety of factors such as the growth stage of the plant, species, and tissue type examined [35]. Therefore this study was complemented by an exploration of the endophytic bacterial species inhabited in the citrus phyllo sphere. Thirty-seven distinct strains of bacteria, including Firmicutes and Proteobacteria, were found. Bacillus sp. was the most prevalent genus observed in citrus.

Endophytes have been identified from approximately all kinds of Vegetation [6, 36, 37]. Numerous studies have also reported the isolation of indigenous endophytes from other hosts, like potatoes [25], maize [38], wheat [39], rice, banana, carrot, sugarcane, tomatoes [28], and peppers [40], soybean plants and among others [41]. To the best of our knowledge, this is the first study to characterize indigenous bacterial endophytes extracted from different citrus varieties.

This remains a substantial difference in the kinds of indigenous microorganisms extracted from distinct different host plants. Bacterial isolates are distinguished by their colors and colony shape [42, 43]. A previous study was revealed the identification of BE from tomato [44], common bean (*Phaseolus vulgaris*) [19], and Alien Weed (*Mikania micrantha*) [21]. Endophytes have also been isolated from the roots, stems, and leaves of numerous rice varieties [45].

A variety of bacteria have already been identified from the vascular tissue of lemon roots, such as *Alcaligenes-Moraxella, Acinetobacter baumanii, Acromobacter spp.,Bacillus spp., Acinetobacter iwoffii, Arthrobacter spp., Corynebacterium spp., Burkholderia cepacia, Citrobacter freundii* [46]. Endophytes such as (*Citrobacter freundii, Achromobacter, Acinetobacter, Arthrobacter, Burkholderia cepacia, Alcaligenes–Moraxella, Enterobacter, Bacillus, Corynebacterium, and Pseudomonas*) have also been found in the vascular tissue of lemon roots in Florida [47]. Endophytic bacteria *Pantoea agglomerans* and *Bacillus pumilus* were also identified from citrus rootstocks from Brazil reported by [48].

### Molecular characterization of bacterial endophytes

The search for novel bacterial endophytes may contribute to the discovery of novel processes that boost plant growth and disclose interesting interactions between plants and bacterial endophytes. Though the use of 16S rDNA sequencing to identify bacterial diversity has resulted in the recognition and characterization of many previously unknown bacteria, determining pathogenic microbes and correlating them to disease still requires bacterial cultures. [49] Conducted similar studies on the 16S rRNA from the soya bean. However, in beans, 16S rRNA sequencing was performed, and a taxonomic study was applied to determine the evolutionary pattern between bacterial strains [50].

### Signature Sequences

Signature patterns of all isolated genera were identified as signature patterns are critical characteristics for recognizing bacteria. There have been few studies to identify the distinctive signatures for another bacterial genus. The signature patterns of 15 genera were detected in this study, both at the species level levels. Every bacteria has a distinctive pattern that is genetically present in the 16S rRNA region. This Unique Pattern may perhaps be advantageous in the diagnosis of specific bacterial strains and also help to determine the biodiversity of bacteria in different environmental samples. By comparing all of the isolates to other reported strains in the database, signatures at the genus and species level were discovered in this study. Target-specific patterns could be determined based on dinucleotide composition to discriminate one group of bacteria from another [51]. These dinucleotides could be used to construct distinct sequence patterns to determine specificity against standard databases. However, the repeating patterns are conserved across different sequences of Pseudomonas that have been used to pinpoint a mismatched region and signature. In this research signatures of Bacillus, Lysinibacillus, Enterobacter, Pseudomonas, Enterococcus, Brevibacterium, and some other genera have been used to identify genus-specific patterns and primers. The 4 detected repeats show a pattern of repeating parts that are highly prevalent across the pseudomonas 16S sequences. At the genus level, the signature sequence revealed a repetition in a sequence because it was not the same from species, which indicated just a single unique sequence could be distinct from other reported species of the same genus. No repeats of the sequence were discovered in certain taxa, as mentioned in (Table 4).

Except for Pseudomonas, the incidence of four repeats in a row was not seen in some other groups. This indicates that in other genera, these patterns have undergone substitutions/changes at one or more corresponding base locations, resulting in repeats not being formed, or that these four conserved repeating patterns in Pseudomonas are the consequence of evolution. Previously finding of identical repeat units throughout vertebrate species, some studies have proposed the evolutionary importance of repeat elements (Wilkinson et al. 1997).

Furthermore, in Pseudomonas, the discovered consistent repeats may have some relevance that should be examined. Another type of positional identifiers, such as non-duplicating patterns, requires the same distances between such sequence patterns. Although the approach was effective for Pseudomonas, it may have limitations when applied to other genera. In other cases, for instance, subsequences bordered by regular repetitions might indicate reduced variation throughout the length of the fragment or the variation may be uniform during its length; in certain cases, identifying a stretch to select patterns would be difficult. Furthermore, it’s indeed possible that the patterns belonging to the most variable region do not produce enough hits to be considered target-specific. It’s unclear to what extent various bacterial endophyte populations can be found in different plant tissues and species. Their characteristics and functional responsibilities, on the other hand, can vary. The hunt for novel bacterial endophytes, on the other hand, may aid in the discovery of novel mechanisms that enhance plant growth, as well as disclose intriguing interactions between plants and their endophytes, as well as between endophyte strains of the plant.

## ACKNOWLEDGMENTS

The author acknowledges Prof. Dr. Muhammad Saleem Haider Dean Faculty of Agriculture Sciences, University of the Punjab, and Lahore for providing research facilities. Special thanks to Dr. Muhammad Shafiq to provide guidance.

## REFERENCES

1. Nawaz MA, Ahmad W, Ahmad S, and Khan MM, et al. Role of growth regulators on pre harvest fruit drop, yield and quality in Kinnow mandarin. Pakistan Journal of Botany.2008; 40(5):1971–1981.

2. Adenaike O, Abakpa GO. Antioxidant Compounds and Health Benefits of Citrus Fruits. European Journal of Nutrition and Food Safety.2021; 65–74.

3. Memon NA. Market potential for Pakistani citrus fruits (Kinnow) in world. Pakistan Food Journal.2014; 1:41–42.

4. Monier JM, Lindow S. Frequency, size, and localization of bacterial aggregates on bean leaf surfaces. Applied and Environmental Microbiology. 2004;70(1): 346–55.

5. Mina D, Pereira JA, Lino-Neto T, Baptista P, et al. Epiphytic and endophytic bacteria on olive tree phyllosphere: exploring tissue and cultivar effect. Microbial ecology.2020; 1–13.

6. Ryan RP, Germaine K, Franks A, Ryan DJ, Dowling DN, et al. Bacterial endophytes: recent developments and applications. FEMS microbiology letters.2008; 278(1):1–9.

7. Papik J, Folkmanova M, Polivkova M, Suman J, Uhlik O, et al. The invisible life inside plants: Deciphering the riddles of endophytic bacterial diversity. Biotechnology advances.2020;107614.

8. Rasche F, Velvis H, Zachow C, Berg G, Van Elsas JD, Sessitsch A, et al. Impact of transgenic potatoes expressing anti-bacterial agents on bacterial endophytes is comparable with the effects of plant genotype, soil type and pathogen infection. Journal of Applied Ecology.2006; 43(3): 555–566.

9. Saleem B. Phyllosphere Microbiome: Plant Defense Strategies. In Microbiomes and the Global Climate Change (pp. 173–201). Springer, Singapore.2021.

10. Schulz B, Boyle C. What are endophytes? Microbial Root Endophytes (Schulz BJE, Boyle CJC & Sieber TN, eds), pp. 1–13. Springer-Verlag, Berlin.2006.

11. Bosamia TC, Barbadikar KM, Modi A, et al. Genomic insights of plant endophyte interaction: prospective and impact on plant fitness. In Microbial Endophytes (pp. 227–249). Woodhead Publishing.2020.

12. Garbeva P, Van Overbeek LS, Van Vuurde JWL, Van Elsas JD, et al. Analysis of endophytic bacterial communities of potato by plating and denaturing gradient gel electrophoresis (DGGE) of 16S rDNA based PCR fragments. Microbial ecology.2001; 41(4): 369–83.

13. Hernández-Pacheco CE, del Carmen Orozco-Mosqueda M, Flores A, Valencia-Cantero E, Santoyo G, et al. Tissue-specific diversity of bacterial endophytes in Mexican husk tomato plants (Physalis ixocarpa Brot. ex Horm.), and screening for their multiple plant growthpromoting activities. Current Research in Microbial Sciences.2021;2:100028.

14. Van Overbeek L, Van Elsas JD. Effects of plant genotype and growth stage on the structure of bacterial communities associated with potato (Solanum tuberosum L.). FEMS Microbiology and Ecology.2008; 64(2):283–96.

15. Compant S, Duffy B, Nowak J, Clément C, Barka EA, et al. Use of plant growth-promoting bacteria for biocontrol of plant diseases: principles, mechanisms of action, and future prospects. Applied and environmental microbiology. 2005;71(9):4951–59.

16. Lamb TG, Tonkyn DW, Kluepfel DA, et al. Movement of Pseudomonas aureofaciens from the rhizosphere to aerial plant tissue. Canadian Journal of Microbiology.1996; 42(11):1112–20.

17. Hallmann J, Quadt-Hallmann A, Mahaffee WF, Kloepper JW, et al. Bacterial endophytes in agricultural crops. Canadian Journal of Microbiology.1997;43:895–14.

18. Chi F, Shen SH, Cheng HP, Jing YX, Yanni YG, Dazzo FB, et al. Ascending migration of endophytic rhizobia, from roots to leaves, inside rice plants and assessment of benefits to rice growth physiology. Applied and environmental microbiology.2005;71(11):7271–7278.

19. Rosenblueth M, Martínez-Romero E. Bacterial endophytes and their interactions with hosts. Molecular plant-microbe interactions.2006;19(8):827–837.

20. Elavazhagan T, Jayakumar S, Balakrishnan V, Chitravadivu C, et al. Isolation of endophytic bacteria from the invasive alien weed, Mikania micrantha and their molecular characterization. American-Eurasian Journal of Scientific Research.2009; 4:154–158.

21. Xin G, Glawe D, Doty SL, et al. Characterization of three endophytic, indole-3-acetic acid-producing yeasts occurring in Populous trees. Mycological Research. 2009;113(9):973–80.

22. West ER, Cother EJ, Steel CC, Ash GJ, et al. The characterization and diversity of bacterial endophytes of grapevine. Canadian Journal of Microbiology.2010; 56(3):209–16.

23. Doty SL, Dosher MR, Singleton GL, Moore AL, Van Aken B, Stettler RF, Gordon MP, et al.Identification of an endophytic Rhizobium in stems of Populus. Symbiosis.2005; 39(1): 27–35.

24. Khan Z, Doty SL.(2009). Characterization of bacterial endophytes of sweet potato plants. Plant and soil.2009; 322(1):197–207.

25. Andreote FD, da Rocha UN, Araújo WL, Azevedo JL, van overbeek LS, et al. Effect of bacterial inoculation, plant genotype and developmental stage on root-associated and endophytic bacterial communities in potato (Solanum tuberosum). Antonie van Leeuwenhoek.2010; 97: 389–99.

26. Manter DK, Delgado JA, Holm DG, Stong RA, et al. Pyrosequencing reveals a highly diverse and cultivar-specific bacterial endophyte community in potato roots. Microbial Ecology.2010; 60(1):157–66.

27. Hung PQ, Kumar SM, Govindsamy V, Annapurna K, et al. Isolation and characterization of endophytic bacteria from wild and cultivated soybean varieties. Bioloy and Fertile Soil.2007;44(1):155–62.

28. Marquez-Santacruz HA, Hernandez-Leon R, Orozco-Mosqueda MC, Velazquez-Sepulveda I, Santoyo G, et al. Diversity of bacterial endophytes in roots of Mexican husk tomato plants(Physalisixocarpa) and their detection in the rhizosphere. Genetic and Molecular Research.2010;9(4):2372–80.

29. Garrity G. The Proteobacteria. Bergeys’s Manaul of Systematic Bacteriology. Springer, New York.2005.

30. Wilson K. Preparation of genomic DNA from bacteria. Current protocols in molecular biology.1987;2–4.

31. Tamura K, Peterson D, Peterson N, Stecher G, Nei M, Kumar S, et al. MEGA5: molecular evolutionary genetics analysis using maximum likelihood, evolutionary distance, and maximum parsimony methods. Molecular Biology and Evology.2011;28(10):2731–2739.

32. Krieg NR, Holt JG. Bergey’s manual of systematic bacteriology (No. BOOK). Yi Hsien Publishing Co.1984.

33. Costa LEO, Queiroz MV, Borges AC, Moraes CA, Araújo EF, et al.(2012) Isolation and characterization of endophytic bacteria isolated from the leaves of the common bean (P. vulharis). Brazilian Journal of Microbiology.2012;43:1562–1575.

34. Trivedi, P., Mattupalli, C., Eversole, K., & Leach, J. E. (2021). Enabling sustainable agriculture through understanding and enhancement of microbiomes. New Phytologist, 230(6), 2129–2147.

35. Santoyo G, Moreno-Hagelsieb G, del Carmen Orozco-Mosqueda M,Glick BR, et al. Plant growth-promoting bacterial endophytes. Microbiological Research.2016;183: 92–99.

36. Dudeja SS, Suneja-Madan P, Paul M, Maheswari R, Kothe E, et al. Bacterial endophytes: Molecular interactions with their hosts. Journal of Basic Microbiology.2021; 61(6): 475–505.

37. Munir S, Li Y, He P, Huang M, He P, He P, He Y, et al. Core endophyte communities of different citrus varieties from citrus growing regions in China. Scientific reports.2020;10(1):1–12.

38. Stamford TLM, Stamford NP, Coelho LCBB, Araujo JM, et al. Production and characterization of a thermostable glucoamylase from Streptosporangium sp. endophyte of maize leaves. Bioresource Technology.2002;83(2):105–109.

39. Majeed A, Abbasi MK, Hameed S, Imran A, Rahim N, et al. Isolation and characterization of plant growth-promoting rhizobacteria from wheat rhizosphere and their effect on plant growth promotion. Frontior Microbiology.2015: 6.

40. Marasco R, Rolli E, Ettoumi B, Vigani G, Mapelli F, Borin S, Zocchi G, et al. A drought resistance-promoting microbiome is selected by root system under desert farming. PloS one.2012;7(10): 48479.

41. Sturz AV, Christie BR, Nowak J, et al. Bacterial endophytes: potential role in developing sustainable systems of crop production. Critical reviews in plant sciences.2000;19(1):1–30.

42. Datta J, Kopczyńska P. Effect of kenaf fibre modification on morphology and mechanical properties of thermoplastic polyurethane materials. Industrial Crops and Products. 2015;74:566–576.

43. Silaban S, Marika DB, Simorangkir M, et al. Isolation and characterization of amylase-producing amylolytic bacteria from rice soil samples. In Journal of Physics: Conference Series (Vol. 1485, No. 1, p. 012006). IOP Publishing.2020.

44. Agrawal DPK, Agrawal S. Characterization of Bacillus sp. strains isolated from rhizosphere of tomato plants (Lycopersicon esculentum) for their use as potential plant growth promoting rhizobacteria. International Journal of Current Microbiology and Applied Scieces.2013;2(10):406–417.

45. Kumar V, Jain L, Jain SK, Chaturvedi S, Kaushal P, et al. Bacterial endophytes of rice (Oryza sativa L.) and their potential for plant growth promotion and antagonistic activities. South African Journal of Botany.2020;134:50–63.

46. Gardner JM, Feldman AW, Zablotowicz RM, et al. Identity and behavior of xylem-residing bacteria in rough lemon roots of Florida citrus trees. Applied and Environmental Microbiology.1982;43(6):1335–1342.

47. Gardner NJ, Savard T, Obermeier P, Caldwell G, Champagne CP, et al. Selection and characterization of mixed starter cultures for lactic acid fermentation of carrot, cabbage, beet and onion vegetable mixtures. International journal of food microbiology, 2001;64(3): 261–275.

48. Araújo WL, Maccheroni W, Azevedo JL, et al. Characterization of an endophytic bacterial community associated with Eucalyptus spp. Genetics and Molecular Research.2009; 8(4):1408–1422.

49. Hung PQ, Annapurna K. Isolation and characterization of endophytic bacteria in soybean (Glycine sp.). Omonrice.2004; 12:92–101.

50. Bhore SJ, Ravichantar N, Loh CY, et al.2010. Screening of endophytic bacteria isolated from leaves of Sambung Nyawa [Gynura procumbens (Lour.) Merr.] for cytokinin-lik compounds. Bioinformation.2010;5(5): 191.

51. More RP, Purohit HJ. The identification of discriminating patterns from 16S rRNA gene to generate signature for bacillus genus. Journal of Computational Biology.2016; 23(8): 651–661.

52. Wilkinson GS, Mayer F, Kerth G, Petri B, et al. Evolution of repeated sequence arrays in the D-loop region of bat mitochondrial DNA. Genetics.1997;146(3): 1035–1048.

